# Phase-amplitude coupling profiles differ in frontal and auditory cortices

**DOI:** 10.1101/2020.05.05.078667

**Authors:** Francisco García-Rosales, Luciana López-Jury, Eugenia González-Palomares, Yuranny Cabral-Calderin, Manfred Kössl, Julio C. Hechavarria

## Abstract

Neural oscillations are at the core of important computations in the mammalian brain. Interactions between oscillatory activities in different frequency bands, such as delta (1-4 Hz), theta (4-8 Hz), or gamma (>30 Hz), are a powerful mechanism for binding fundamentally distinct spatiotemporal scales of neural processing. Phase-amplitude coupling (PAC) is one such plausible and well-described interaction, but much is yet to be uncovered regarding how PAC dynamics contribute to sensory representations. In particular, although PAC appears to have a major role in audition, the characteristics of coupling profiles in sensory and integration (i.e. frontal) cortical areas remain obscure. Here, we address this question by studying PAC dynamics in the frontal-auditory field (FAF; an auditory area in the bat frontal cortex) and the auditory cortex (AC) of the bat *Carollia perspicillata*. By means of simultaneous electrophysiological recordings in frontal and auditory cortices examining local-field potentials (LFPs), we show that the amplitude of gamma-band activity couples with the phase of low-frequency LFPs in both structures. Our results demonstrate that the coupling in FAF occurs most prominently in delta/high-gamma frequencies (1-4/75-100 Hz), whereas in the AC the coupling is strongest in the theta/low-gamma (2-8/25-55 Hz) range. We argue that distinct PAC profiles may represent different mechanisms for neuronal processing in frontal and auditory cortices, and might complement oscillatory interactions for sensory processing in the frontal-auditory cortex network.

## Introduction

There is increasing evidence supporting the role of oscillatory activity as instrument of neural computations in the mammalian brain. Oscillations in low- and high-frequencies, particularly in the delta-to-alpha (1-12 Hz) and gamma (>30 Hz) ranges, are deemed essential for numerous tasks, including sensory processing and selectivity (Schroeder & Lakatos, 2009; Bosman *et al.*, 2012; Obleser & Kayser, 2019), the implementation of attentional mechanisms (Lakatos *et al.*, 2013; Magazzini & Singh, 2018), cognitive control (Cho *et al.*, 2006; Helfrich & Knight, 2016), learning and memory (Benchenane *et al.*, 2010; Wang *et al.*, 2018), or inter-areal connectivity by means of communication-through-coherence (Fries, 2015). High- and low-frequency oscillations represent the activity of local and global neuronal ensembles, respectively, occurring at different timescales determined by the oscillatory frequencies (Canolty & Knight, 2010). The question of how different spatiotemporal scales are integrated in the brain, and therefore the relationship between co-existing low- and high-frequency activities, has gained attention in recent years.

Cross-frequency coupling is a plausible mechanism that could allow for the binding of low- and high-frequency oscillations and their respective spatiotemporal dynamics (Canolty & Knight, 2010; Tort *et al.*, 2010; Hyafil *et al.*, 2015b). A specific form of cross-frequency coupling, namely phase-amplitude coupling (PAC), has been related to numerous brain functions. PAC is the phenomenon whereby the phase of a low-frequency oscillation couples with the amplitude of a high-frequency one. This type of interaction between distinct frequency bands is well established in regions such as the hippocampus (Lisman & Jensen, 2013), the frontal cortex (Helfrich & Knight, 2016), and cortical sensory areas (Spaak *et al.*, 2012; Esghaei *et al.*, 2015; O’Connell *et al.*, 2015; Sotero *et al.*, 2015; Xiao *et al.*, 2019). PAC in these regions has been associated with working memory (Axmacher *et al.*, 2010; Daume *et al.*, 2017), learning (Tort *et al.*, 2009), behavioural coordination (Amemiya & Redish, 2018), and the organization of inter-areal communication and information binding (Colgin *et al.*, 2009; Daume *et al.*, 2017). High-order sensory processing may also capitalize on PAC, the latter providing a mechanistic substrate for the parsing of continuous stimuli by accommodating local network activity in the gamma range into slower, behaviourally relevant timescales represented by the low-frequency activity (Giraud & Poeppel, 2012; Hyafil *et al.*, 2015b). Indeed, theta-gamma coupling in the auditory cortex (AC) of humans has been suggested as vital component of speech processing (Giraud & Poeppel, 2012; Morillon *et al.*, 2012; Gross *et al.*, 2013; Hyafil *et al.*, 2015a), while it has been shown that similar PAC profiles in the primate AC mediate acoustic sequence learning (Kikuchi *et al.*, 2017).

At present, little is known about PAC dynamics in auditory regions of animal models beyond primates. However, tackling such question can provide valuable insights into the nature of evolutionarily preserved circuits across species. In both primates and non-primates there exist structures in the frontal cortex that are strongly responsive to acoustic stimuli (Kobler *et al.*, 1987; Eiermann & Esser, 2000; Medalla & Barbas, 2014; Plakke & Romanski, 2014). Within these structures, the relationship between the phase of low frequency oscillations and the amplitude of high frequency rhythms remains largely unexplored. The current study aims to bridge this gap by means of electrophysiological recordings of local-field potentials (LFPs) from the AC and a region of the short-tailed bat’s (*Carollia perspicillata*) frontal cortex, specialized for audition: the frontal-auditory field (FAF; (Kobler *et al.*, 1987; Eiermann & Esser, 2000)). In previous work we showed robust oscillatory responses to acoustic stimulation in *C. perspicillata*’s FAF and AC (Hechavarria *et al.*, 2016b; García-Rosales *et al.*, 2020), and that in the latter structure low-frequency LFPs could be crucial for the neuronal coding of naturalistic sequences (García-Rosales *et al.*, 2018a; García-Rosales *et al.*, 2018b). Furthermore, we observed functional coupling in the FAF-AC circuit by means of delta- (without auditory input) and gamma-band LFPs (García-Rosales *et al.*, 2020). The above supports the roles of cortical oscillations for auditory computations in fronto-temporal networks in mammals.

In this paper, we characterized PAC profiles in FAF and AC during spontaneous activity and sound processing. We found that high-frequency amplitude was coupled to the phase of low-frequency rhythms in frontal and auditory cortices. However, the specific frequencies at which this occurred differed across structures, both with and without acoustic stimulation. Delta/high-gamma PAC was typical in FAF, whereas theta/low-gamma coupling occurred most prominently in the AC. We argue that distinct PAC profiles in FAF and AC could represent distinct mechanisms of neural processing at the level of sensory and association areas.

## Results

### Neural responses in the FAF and AC of C. perspicillata

Electrophysiological experiments were performed on 5 adult male *Carollia perspicillata* bats. We recorded a total of 50 penetrations pairs, each pair comprised of simultaneously recorded neural activity in the FAF and AC. In the FAF, a single carbon electrode was inserted, and recordings were made at depths ranging from 300-450 μm. In the AC, a 16-channel probe was used, allowing to record from cortical depths spanning 0-750 μm at once. Both spontaneous (i.e. without acoustic stimulation) and auditory-driven neural activities were analysed. Auditory stimuli consisted of a natural call (henceforth, “nat” stimulus) representative of this species distress repertoire (**Fig. 1A**; see (Hechavarria *et al.*, 2016a)), as well as two artificially constructed syllabic trains (see Methods and (García-Rosales *et al.*, 2020)). The first of these trains had an isochronous structure by which a natural syllable was repeated with a rate of 5.28 Hz (**Fig. 1B**). In the second train, the same syllable was repeated in a Poisson-like manner (i.e. non-periodically) with an average rate of 70 Hz (**Fig. 1C**). The natural syllable used to construct these trains was also typical for distress syllables in this bat species (Hechavarria *et al.*, 2016a); its time-frequency representation is shown in **Fig. 1D**.

**Figure 1.**
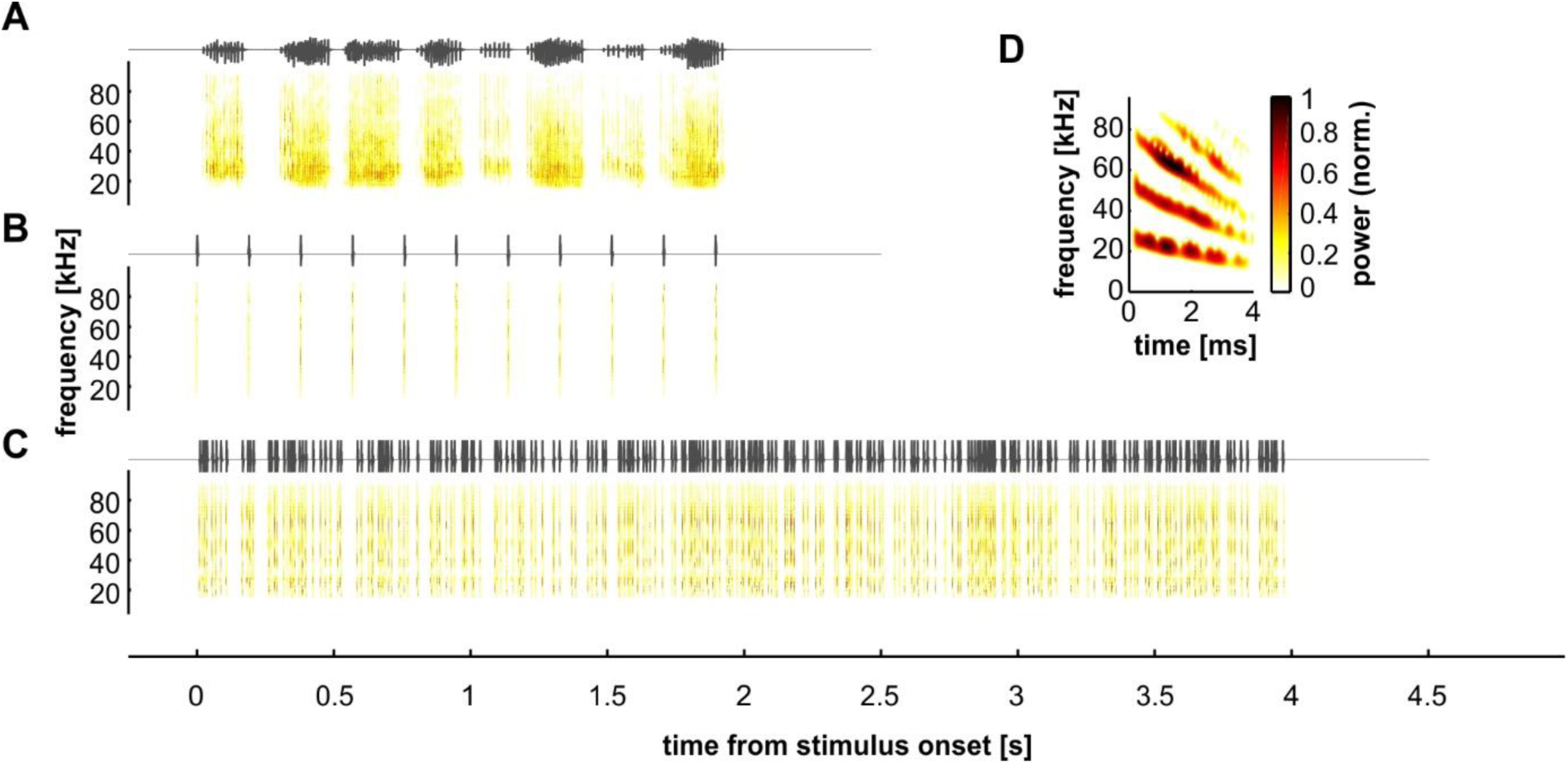
Auditory stimuli. Oscillograms (top) and spectrograms (bottom; normalized amplitude and power) of the sounds used for auditory stimulation. These comprised a natural call (**A**), a syllable train with a repetition rate of 5.28 Hz (**B**), and a syllable train with a Poisson temporal structure (**C**). Panel **D** shows the spectrotemporal design of the natural distress syllable used to construct the trains in **B** and **C**.

Local-field potentials were analysed for each penetration pair in FAF and AC. We observed robust auditory responses in the LFPs of both structures, as illustrated for a representative penetration pair in **Fig. 2A-C**, where single trial responses to each stimulus (coloured traces) are shown together with the trial-average LFPs for the penetration (black). In the panels of **Figure 2**, data are shown from the FAF and the AC, the latter at a depth of 450 μm (top and bottom subpanels, respectively). Single trials in FAF and AC depicted with the same colour were recorded simultaneously. **Figure 2D** illustrates LFP chunks (see Methods) corresponding to spontaneous activity. Note that no trial-average is shown because there are no temporal references for inter-chunk averaging, such as the onset of a stimulus.

**Figure 2.**
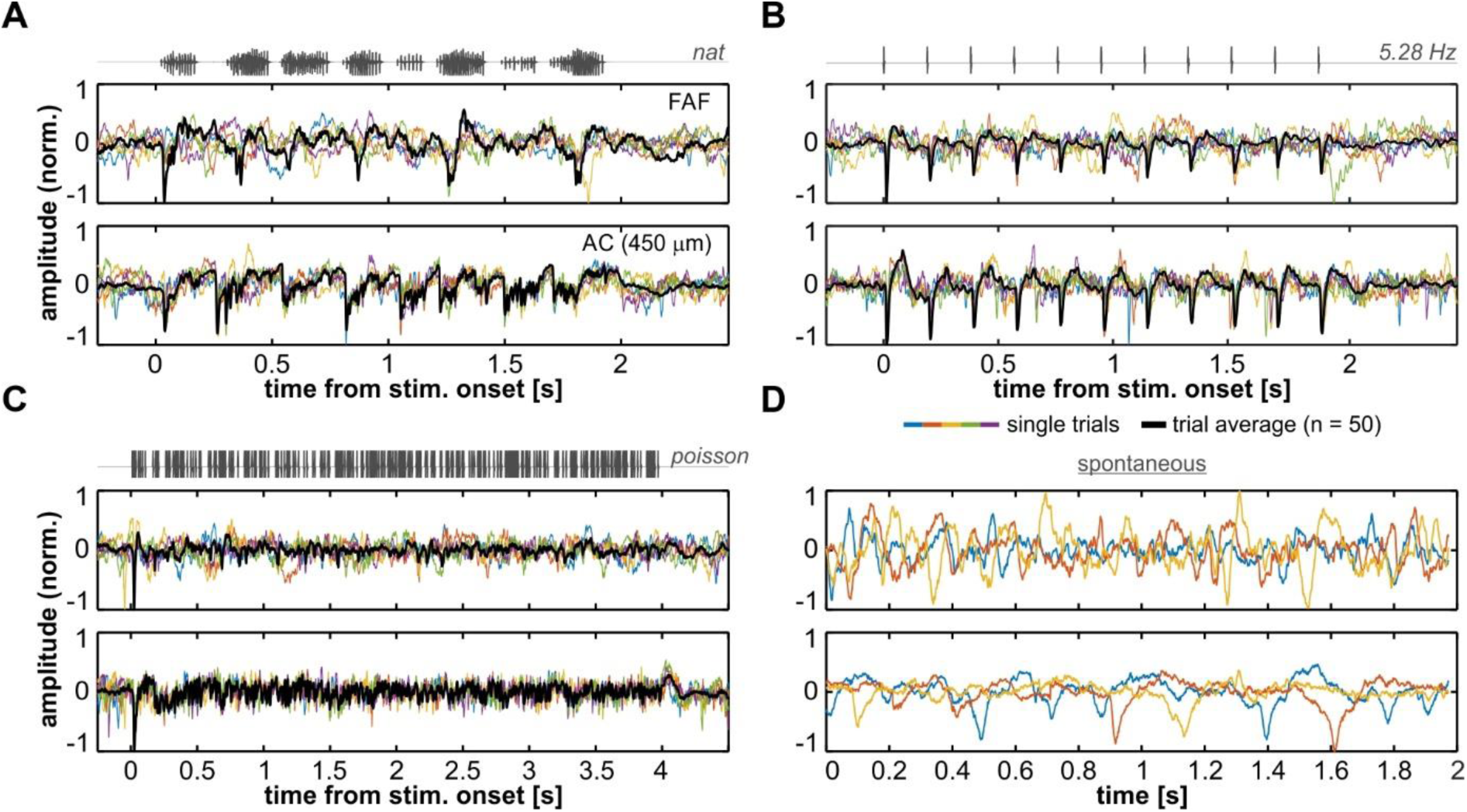
Representative LFP recordings during auditory processing and spontaneous activity. (**A**) LFP traces from the FAF (top) and the AC at a depth of 450 μm (bottom), in response to the natural call (stimulus’ oscillogram shown on top of the subpanels). Coloured lines correspond to five single-trial recordings (out of a total of 50 trials) from a representative penetration. The thick black line depicts the trial average. (**B**) Same as in **A**, but LFPs were recorded in response to the 5.28 Hz syllabic train. (**C**) Same as in **A** and **B**, but responses correspond to the Poisson train. (**D**) Three representative chunks of spontaneously recorded LFP signals of the same penetration shown in **A**-**C**. Note that a trial average is lacking as chunks do not share a reference time point (e.g. stimulus onset). In all panels, single-trial LFP traces from FAF and AC with the same colour were recorded simultaneously.

### Phase amplitude-coupling in FAF and AC

We evaluated phase-amplitude coupling between low and high frequencies in FAF and AC using a procedure inspired by a previous study (Kikuchi *et al.*, 2017). LFPs for phase were filtered (4^th^ order bandpass Butterworth) in frequency bands with centres at 2, 4, 6, …, 14 Hz, and 2 Hz bandwidth. LFPs for amplitude were filtered (same filter as before) in frequency bands centred at 30, 35, 40, …, 125 Hz, with 10 Hz bandwidth. **Figure 3A** illustrates two single-trial recordings from the FAF in response to the Poisson train. Delta- (1-3 Hz) and gamma-band (75-85 Hz; bands chosen for illustrative purposes) LFPs are shown in grey and red, respectively. The instantaneous phase and amplitude of delta and gamma LFP signals are also shown in black and orange, respectively. Instantaneous phase and amplitude were extracted after Hilbert-transforming the filtered LFPs. To correct for possible biases due to phase non-uniformity in the signals (Aru *et al.*, 2015; van Driel *et al.*, 2015), the mean phase vector was linearly subtracted from the instantaneous phase series. PAC in FAF and AC (PAC_FAF_ and PAC_AC_, respectively) were quantified by z-normalizing a modulation index (MI; z-normalized index: zMI) to a surrogate distribution where effects of evoked-related responses across trials were tackled (**Fig. 3A, B**; see Methods for details). Based on the zMIs we obtained PAC profiles for each channel and penetration.

**Figure 3.**
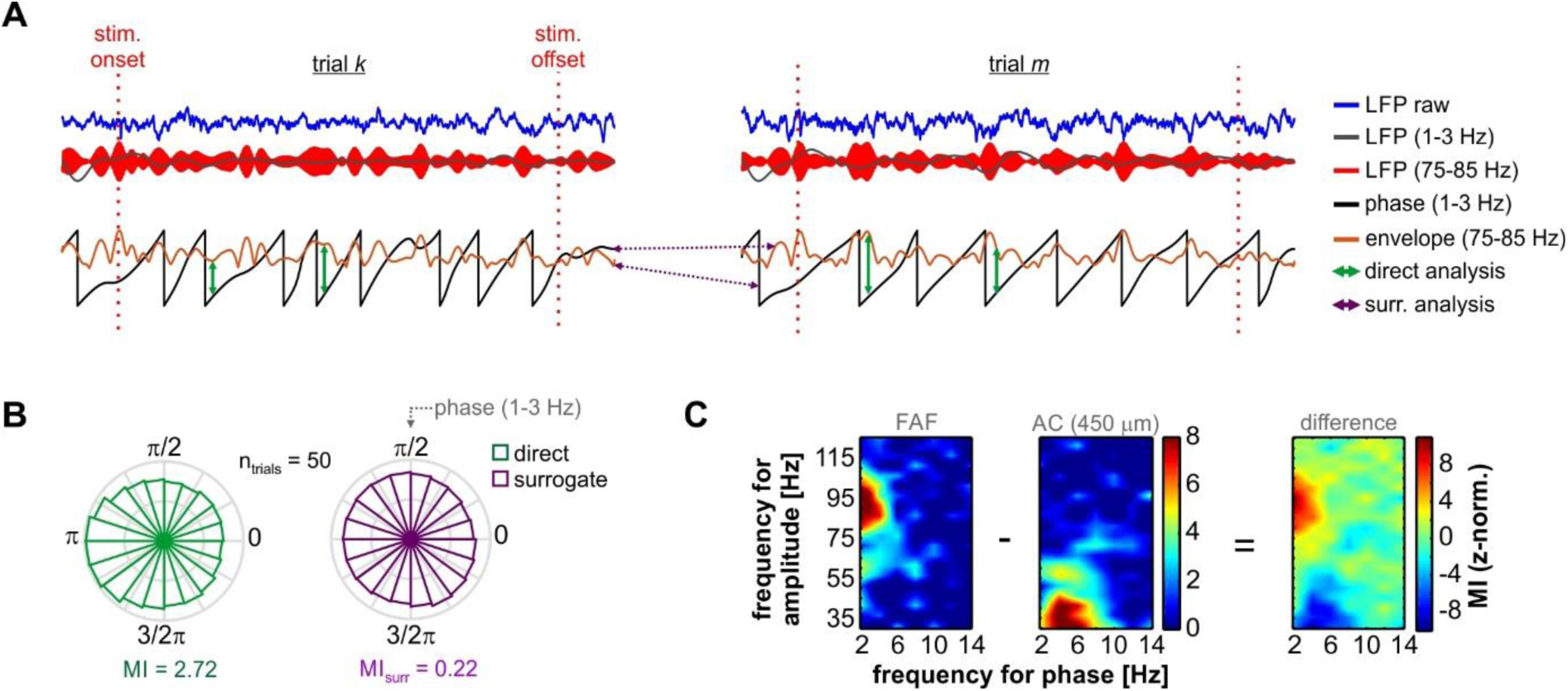
Phase-amplitude coupling analyses in representative recordings. (**A**) Single trial LFPs (2 exemplary trials *k* and *m*, blue traces), recorded from the FAF, in response to the Poisson syllabic train. Stimulus onset and offset are marked with red vertical dashed lines; the window between onset and offset was used for PAC analyses across stimuli (here, 4 s length). Delta- (1-3 Hz) and gamma-band (75-85 Hz) filtered LFPs are depicted in gray and red, respectively. Below, the phase of the delta LFPs (black) and the amplitude envelope of gamma (orange) are shown. In a direct PAC analysis, phase and amplitude were matched within trials (represented with green arrows in the figure). However, during surrogate analyses, amplitude and phase were matched between trials at random (trial shuffling; see Methods). The latter is depicted as purple arrows across trials *k* and *m.* (**B**) Circular distribution depicting the relationship between gamma-band amplitude and delta-band phase. With the direct analysis, the phase-amplitude relationship was visibly non-circularly uniform (green), yielding a modulation index (MI) of 2.72. In an instance of the surrogate analysis (purple), shown for illustrative purposes, the phase-amplitude relationship was distributed rather uniformly, yielding a MI of 0.22. (**C**) Phase-amplitude coupling (PAC) maps calculated from LFPs corresponding to the same penetration shown in **A** and **B**, also in response to the Poisson train. The PAC is shown for the FAF and the AC at a depth of 450 μm. To evaluate differences in the PAC across structures, maps from FAF and AC were subtracted in further analyses (here shown as “difference”).

Across penetrations, and as is readily visible in **Fig. 3C**, the PAC in the FAF typically peaked in the delta/high-gamma range of the PAC maps (δ/γ_high_; i.e. frequency for phase: ∼1-4 Hz, frequency for amplitude: ∼70-100 Hz). On the other hand, the PAC in the AC was strongest in channels located in input layers (depths of 300-550 μm), and the values of zMI typically peaked in the delta-theta/low-gamma range of the maps (θ/γ_low_; frequency for phase: ∼2-8 Hz, frequency for amplitude: ∼25-55 Hz). This trend occurred even without acoustic stimulation, as depicted in **Fig. 4A**, where population-level spontaneous PAC profiles are shown for the FAF and the AC, the latter at representative depths of 50, 300, 450, 600, and 750 μm. We observed that zMIs across the population were significantly higher than 0 in both FAF and AC (regions delimited by grey contour lines; FDR-corrected tailed Wilcoxon signed rank tests, p_corr_ < 0.05), and that significant zMIs within single penetrations (i.e. zMIs > 2.5; see Methods) occurred in ∼25% of the cases (out of n = 49 penetrations; 1 penetration could not be examined during spontaneous activity) in FAF, mostly located in the δ/γ_high_ range of the PAC space, and in ∼40% of the cases in the AC (most strongly for depths of 450 μm), mostly in the θ/γ_low_ range of the PAC space. The quantification of the percentage of significant PAC across the population is shown in **Supplementary Figure 1**.

**Figure 4.**
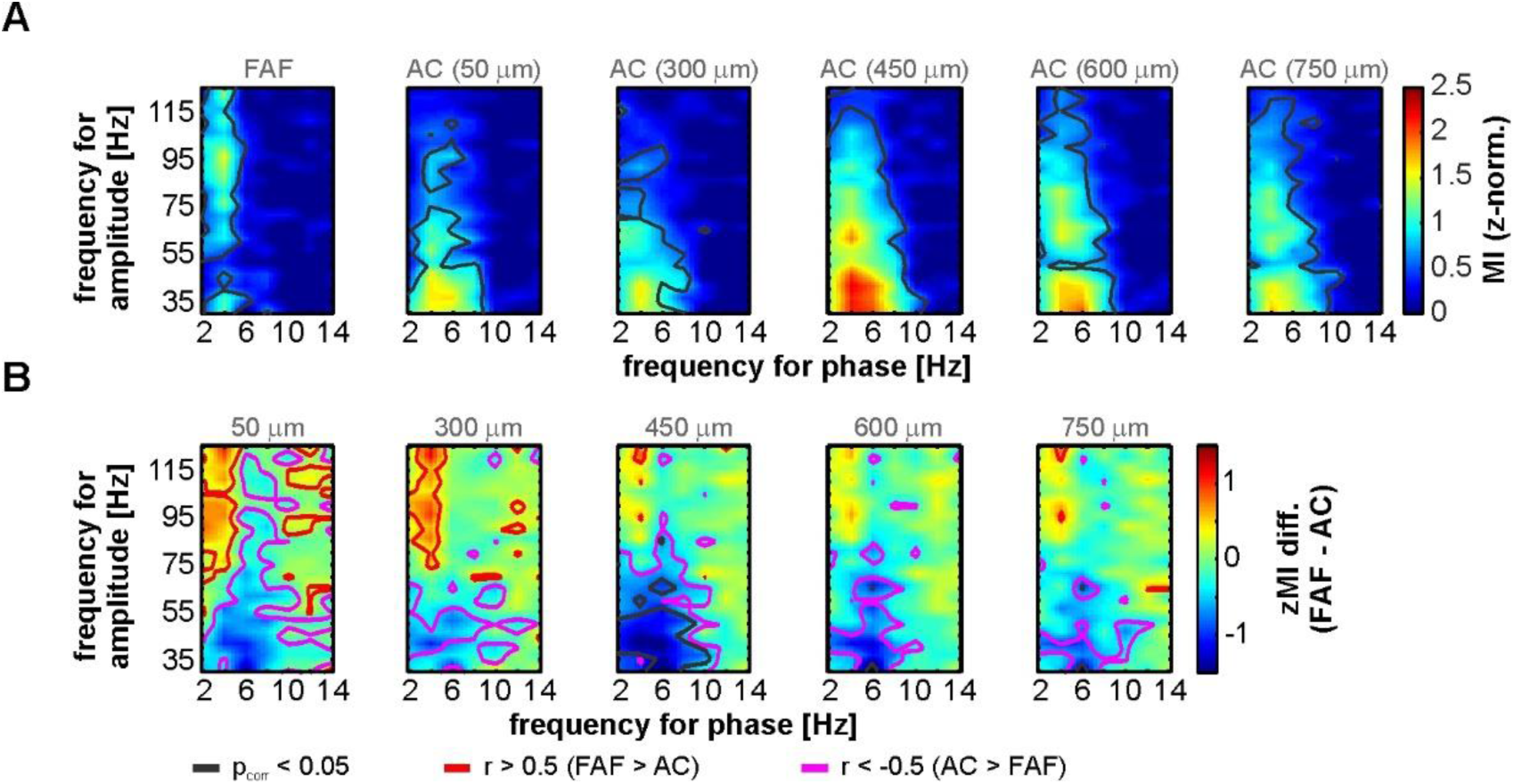
Spontaneous PAC in FAF and AC. (**A**) Population-averaged PAC maps calculated from spontaneously recorded LFPs in the FAF and AC (here shown at depths of 50, 300, 450, 600 and 750 μm). Regions within gray contour lines correspond to those for which z-normalized MIs were significantly above 0 across penetrations (n = 49; FDR-corrected Wilcoxon signed rank test, p_corr_ < 0.05). (**B**) PAC-difference maps between FAF and AC at the same depths from **A**. Red contour lines delimit regions where PAC in FAF (PAC_FAF_) was higher than the PAC in AC (PAC_AC_), with a large effect size (*r* > 0.5). Purple contour lines delimit regions where the opposite occurred (i.e. PAC_AC_ > PAC_FAF_). Gray contour lines mark PAC regions where the differences were significant, after an FDR-corrected Wilcoxon signed rank test (comparing PAC_FAF_ vs PAC_AC_), at an alpha of 0.05.

Regarding the regions in which the coupling peaked, there was a clear distinction between the PAC_FAF_ and the PAC_AC_. This difference was evaluated by subtracting the PAC profiles from the FAF and those from the AC, as illustrated in **Fig. 3C**. By systematically doing so in our dataset during spontaneous activity, differences between PAC_FAF_ and PAC_AC_ became evident (**Fig. 4B**). The spontaneous PAC_FAF_ was higher than the PAC_AC_ in the δ/γ_high_ range, particularly for superficial layers (0-300 μm) of the AC. In this case, although significance did not survive multiple comparisons (FDR-corrected Wilcoxon signed rank tests, p_corr_ > 0.05), effect size estimations (*r*; see Methods) yielded large effects (*r* > 0.5, after (Fritz *et al.*, 2012)) in the δ/γ_high_ range (red contour lines in **Fig. 4B**). Large differences did not occur in this PAC range when considering middle or deep layers of the AC, which could be attributable to the extent of relatively high PAC values to gamma frequencies up to ∼80 Hz (see **Fig. 4A**). In addition, spontaneous PAC_AC_ values were higher than PAC_FAF_ ones in the θ/γ_low_ range, reaching significance at AC depths of 450 μm (grey contour lines in **Fig. 4B**; FDR-corrected Wilcoxon signed rank tests, p_corr_ < 0.05). Still, even in cases where significance did not survive corrections for multiple comparisons (i.e. superficial and deep laminae), we observed large effect sizes indicating that there was a consistent trend of PAC_AC_ being higher than PAC_FAF_ in θ/γ_low_ frequencies (*r* > 0.5, purple contour lines in **Fig. 4B**).

### Population-level PAC in FAF and AC during acoustic processing

The PAC patterns in frontal and auditory cortices remained almost qualitatively unaltered during acoustic stimulation. **Figure 5** depicts PAC maps from AC and FAF, in a similar arrangement to **Fig. 4A**, but using stimulus-driven LFPs recorded in response to the natural call, and to the 5.28 Hz and Poisson syllabic trains (**Fig. 5A-C**, respectively). Again, while population-level PAC_FAF_ was strongest in the δ/γ_high_ range, the PAC_AC_ peaked in the θ/γ_low_ range, more markedly at depths of 450 μm. The former was true independently of the stimulus used. Indeed, we observed that zMIs in FAF and AC were significantly above zero across penetrations (**Fig. 5**, grey contour lines; FDR-corrected tailed Wilcoxon signed rank tests, p_corr_ < 0.05), and that the PAC regions where this happened were predominantly those of δ/γ_high_ and θ/γ_low_ in FAF and AC, respectively. Individually within penetrations and across stimuli, we observed significant zMIs (> 2.5; see Methods) in the FAF occurring in δ/γ_high_ frequencies for ∼40-56% of the penetrations. Significant zMIs were observed in the AC, at 450 μm, for ∼20-25% of the penetrations (see also **Fig. S1**). The decline in the percentage of significant zMIs during acoustic stimulation in the AC can be explained by the stringency of the surrogate analyses used, and by the efforts made to minimize the effect of stimulus-evoked responses in PAC calculations (see Methods). Note that the surrogate analyses, in conjunction with the subtraction of the mean across trials, may obscure PAC values that are not only explained by broadband evoked response, but that are however temporally locked to the stimuli. These did not seem to strongly affect the data from the FAF. The increase of percentage of significant penetrations within FAF in the δ/γ_high_ range indicates a modulation of PAC strength by acoustic stimulation.

**Figure 5.**
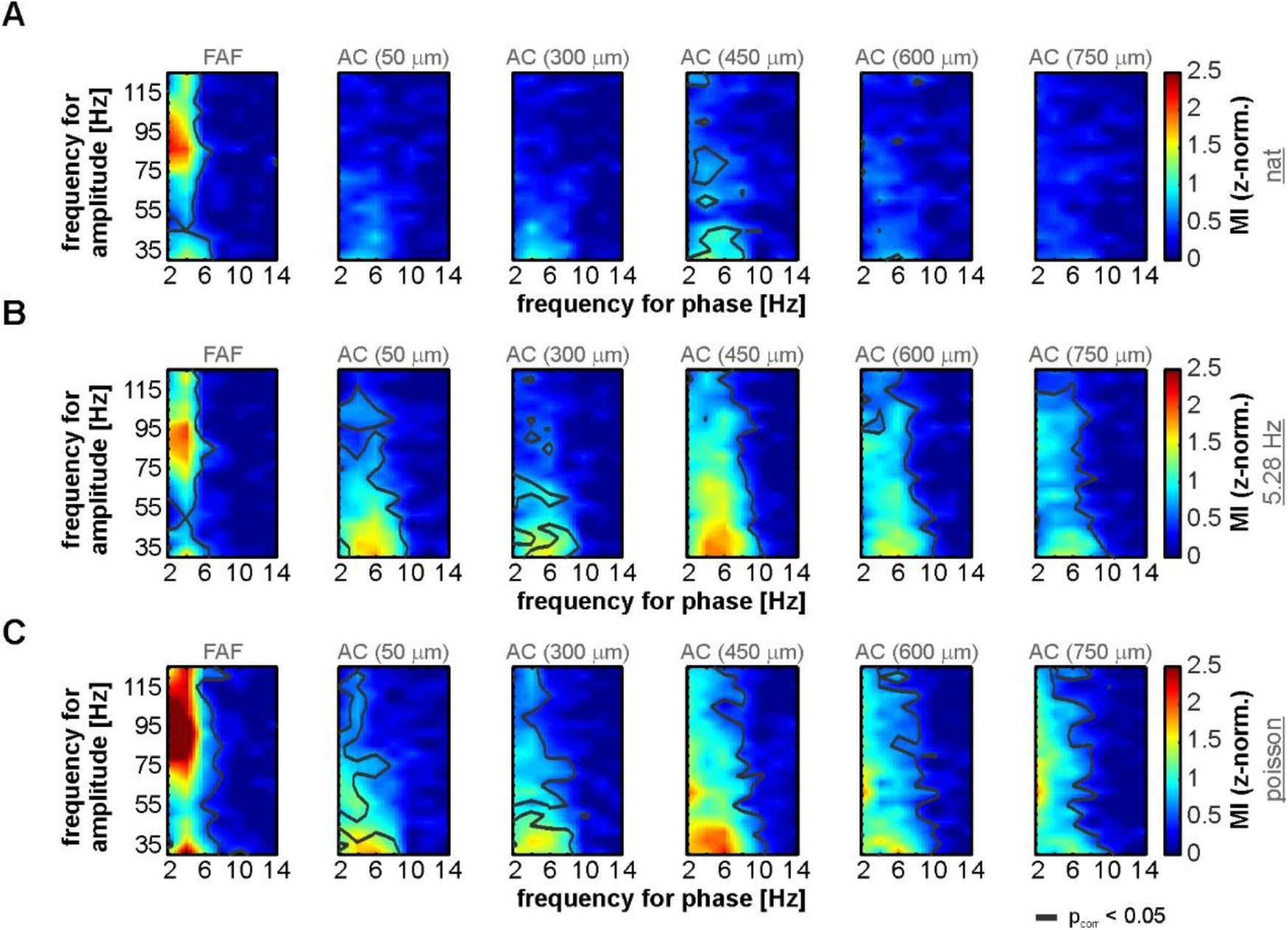
PAC in FAF and AC during acoustic stimulation. (**A**) Population-averaged PAC maps calculated from LFPs recorded in the FAF and the AC at various depths (50, 300, 450, 600, 750 μm), in response to the natural call used as stimulus in this study. Gray contour lines delimit regions where the z-normalized MI was significantly higher than 0 across penetrations (n = 50; FDR-corrected Wilcoxon signed rank test, p_corr_ < 0.05). (**B-C**) Same as in A, but LFPs were recorded in response to the 5.28 Hz (**B**) and the Poisson (**C**) syllabic trains. Note that, independently of the stimulus considered, the PAC in FAF and AC are clearly strongest at distinct phase-amplitude regimes.

Differences between PAC_FAF_ and PAC_AC_ during acoustic processing occurred predominantly in the δ/γ_high_ and θ/γ_low_ ranges (**Fig. 6**). The PAC_FAF_ was significantly stronger than the PAC_AC_ in δ/γ_high_ frequencies at all recording depths of the AC (FDR-corrected Wilcoxon signed rank tests, p_corr_ < 0.05), with large effect sizes (*r* > 0.5; red contour lines in **Fig. 6**) for all stimuli. Conversely, the PAC_AC_ appeared stronger than the PAC_FAF_ in θ/γ_low_ frequencies, although without clear significance for every stimulus tested (statistics as above). In response to the natural call, there were no strong differences between PAC_FAF_ and PAC_AC_ in the θ/γ_low_ range, and effect sizes were also not large (**Fig. 6A**). In the case of the 5.28 Hz (**Fig. 6B**) and the Poisson (**Fig. 6C**) syllabic trains, differences in the PAC across structures were stronger, reaching significance (p_corr_ < 0.05; gray contour lines) mostly at a depth of 450 μm for θ/γ_low_ frequencies. Although for this frequency range significant differences between FAF and AC did not occur with PAC values calculated in response to the Poisson train (compare **Fig. 6B** and **Fig. 6C**, depicting PAC differences for the 5.28 Hz and the Poisson stimuli), large effect sizes were still observed for θ/γ_low_ frequencies (purple contour lines in **Fig. 6** marking areas where PAC_AC_ > PAC_FAF_). Overall, these results indicate that the FAF and the AC engage in distinct phase-amplitude coupling dynamics comprising delta, theta, low- and high-gamma bands of the LFP.

**Figure 6.**
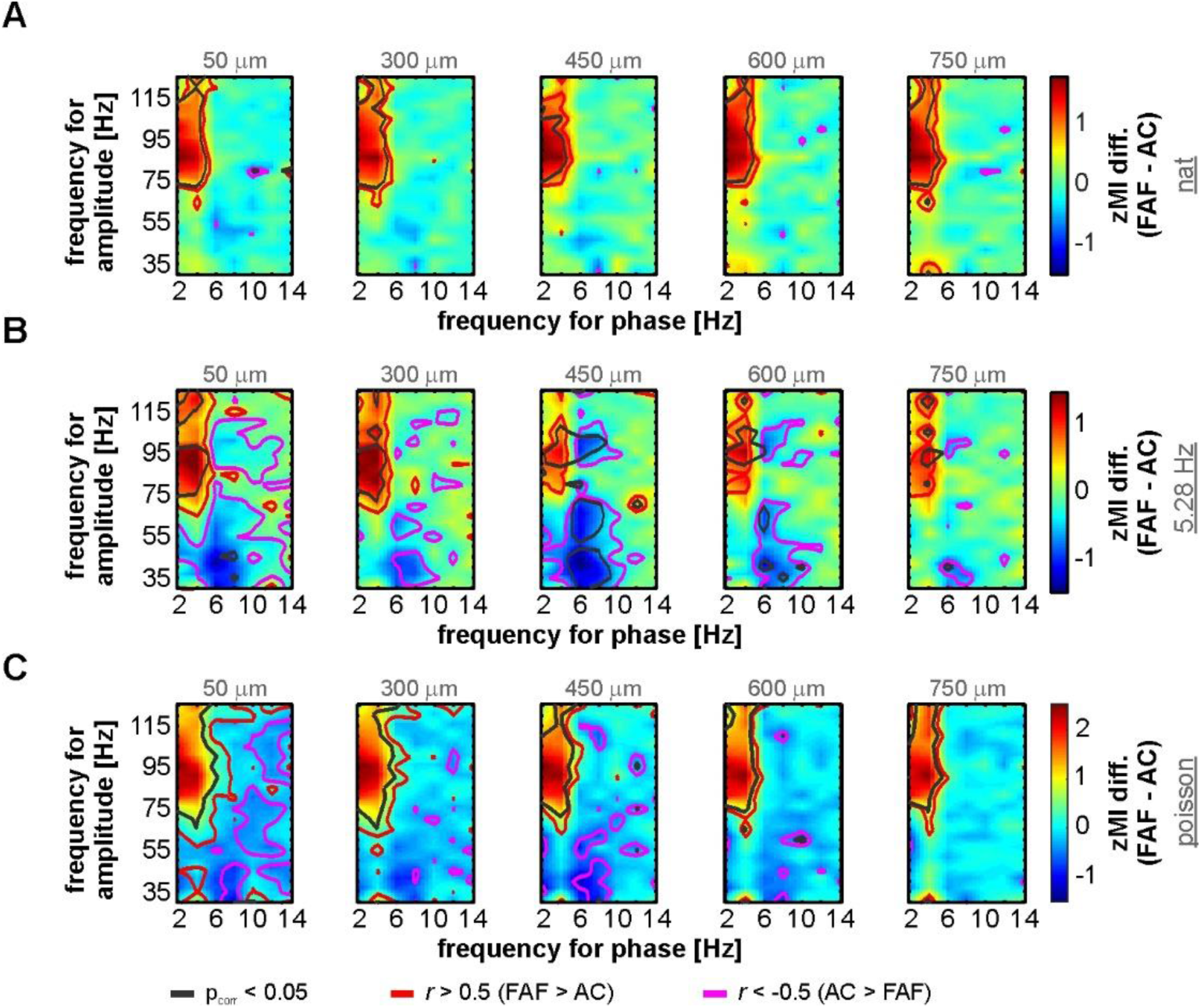
Differences in PAC from FAF and AC during acoustic stimulation. (**A**) Population-averaged difference maps (PAC_FAF_ - PAC_AC_, at various depths in the AC: 50, 300, 450, 600, 750 μm; n = 50) obtained from LFP responses to the natural called used in this study. Red contour lines delimit regions where PAC_FAF_ > PAC_AC_, with large effect sizes (*r* > 0.5), whereas purple contour lines demarcate regions where the opposite occurred (i.e. PAC_AC_ > PAC_FAF_), also with large effect sizes. Gray contour lines mark regions wherein the z-normalized MIs in FAF and AC were significantly different, according to a FDR-corrected Wilcoxon signed rank test, with an alpha of 0.05. (**B-C**) Same as in **A**, but corresponding to LFPs recorded in response to the 5.28 Hz (**B**) and the Poisson (**C**) syllabic trains.

## Discussion

This study addressed the phase-amplitude coupling of oscillatory activities in the AC and FAF of the bat *C. perspicillata*. We report significant PAC in both structures during spontaneous activity and acoustic processing. However, the coupling between low- and high-frequency LFPs differed in auditory and frontal regions of the brain: in the AC, the PAC was strongest in the theta/low-gamma range, while in the FAF the PAC peak occurred predominantly in delta/high-gamma frequencies. Thus, we show that *C. perspicillata*’s AC and FAF exhibit distinct phase-amplitude coupling profiles.

### Phase-amplitude coupling in auditory and frontal areas

Phase-amplitude coupling could be important for the integration of distinct spatiotemporal dynamics represented via low- and high-frequency oscillations (Canolty & Knight, 2010; Hyafil *et al.*, 2015b). Thus, the modulation of high-frequency amplitude by low-frequency phase constitutes a plausible and powerful mechanism of sensory processing. A large body of evidence indicates that slow oscillatory activity in sensory cortices entrains (i.e. synchronizes) to the temporal structure of external stimuli (Sieben *et al.*, 2013; Brookshire *et al.*, 2017; García-Rosales *et al.*, 2018a; Molinaro & Lizarazu, 2018; Doelling *et al.*, 2019).

Synchronized oscillatory activity can act as a sensory filtering mechanism (otherwise called “sensory selectivity”) that is in turn susceptible to top-down modulation by processes such as attention (Lakatos *et al.*, 2008; Schroeder & Lakatos, 2009; Calderone *et al.*, 2014; Obleser & Kayser, 2019). Low frequency activity, coupled with gamma-band oscillations, could therefore align enhanced processing periods (marked by the gamma rhythms) to the structure of external inputs or internal states, thereby boosting their representation in the brain. Research in the human auditory cortex, for example, suggests that theta-gamma PAC plays a major role for the efficient processing of speech signals (Gross *et al.*, 2013; Zion Golumbic *et al.*, 2013; Lizarazu *et al.*, 2019). Indeed, computational modelling demonstrates that the low-frequency stimulus-related neural oscillations might provide a temporal reference frame in which syllables can be processed by high-frequency activity coupled to the underlying slow rhythm (Hyafil *et al.*, 2015a).

It is worth noticing that the effects of PAC in the auditory cortex generalize beyond speech processing. The coupling of low- and high-frequency oscillations could underpin the segmentation of continuous stimuli into behaviourally relevant “perceptual” units. Remarkably, the formation of perceptual auditory units could be particularly important for echolocating bats such as *C. perspicilllata*. A bat exploring its surroundings using sonar receives a stream of incoming echoes corresponding to self-emitted echolocation pulses that reflect from external objects. The integration of those echoes at the level of the auditory or frontal cortices may allow the bat to form an acoustic picture of the environment. Such integration may find its mechanistic substrate in PAC interactions between oscillations working at slow, integrative timescales (e.g. delta, theta, or alpha) and faster ones (e.g. gamma) that would encode the fine structure of the echo streams. Considering that *C. perspicillata* produces echolocation calls with a rate of 40-50 Hz (Beetz *et al.*, 2019), and that LFPs in this bat’s AC strongly synchronize to fast acoustic inputs also in the gamma range (Hechavarria *et al.*, 2016b; Garcia-Rosales *et al.*, 2019), the above-discussed possibility constitutes an interesting view that still requires empirical validation.

In general, it is plausible that PAC dynamics in sensory systems could complement the well-described roles of low-frequency activity for high-order sensory processing (Arnal *et al.*, 2015). Hence, comparable PAC-related phenomena may occur not only across primate species, but also in other mammals. Considering that in the AC of *C. perspicillata* the neuronal representation of acoustic stimuli shares coding mechanisms similar to those described in the primate auditory and visual domains (Belitski *et al.*, 2008; Kayser *et al.*, 2009; Belitski *et al.*, 2010; García-Rosales *et al.*, 2018a; García-Rosales *et al.*, 2018b), the theta-gamma PAC reported in this study could also reflect a general mechanism shared across species. The spontaneous θ/γ_low_ coupling in *C. perspicillata*’s AC and FAF further suggests that the relationship between slow and fast oscillations echoes the properties of cortical networks that do not directly depend on sensory inputs, but that can nevertheless be affected or modulated by them. We therefore propose that auditory cortical theta/low-gamma coupling might provide in bats the same functional advantages proposed for other animals. These circuit dynamics might be evolutionarily preserved, being similar in phylogenetically distant species such as bats and primates.

The frontal cortex is considered an association area where sensory stimuli are integrated, and behavioural/cognitive functions controlled (Miller, 2000; Sugihara *et al.*, 2006; Hage & Nieder, 2015; Carlen, 2017; Hardung *et al.*, 2017). These processes may be supported by oscillatory activity in frontal regions, which further allows coordination with distant areas in the brain including sensory systems and the hippocampus (Park *et al.*, 2015; Helfrich & Knight, 2016; Daume *et al.*, 2017). The frontal-auditory field, located in the bat frontal cortex (Kobler *et al.*, 1987; Eiermann & Esser, 2000), receives auditory inputs from the AC and via a non-lemniscal pathway directly from the suprageniculate nucleus of the thalamus, bypassing major centres such as the inferior colliculus an the AC itself (Kobler *et al.*, 1987; Casseday *et al.*, 1989). The FAF is thus in a privileged position to integrate relatively “raw” auditory information arriving from the thalamus and arguably more processed inputs from the AC. It is likely that the response properties of the FAF, potentially explained by slow afferent synaptic dynamics (Lopez-Jury *et al.*, 2019), constitute evidence for auditory integration in the frontal cortex. This integration can capitalize on PAC, combining low frequency oscillations, which could relate to integratory timescales, with high frequency activity, in turn marking local computations. Although the roles of the FAF for auditory-guided behaviour are not wholly clear, there is evidence indicating that oscillations in this region could coordinate interareal communication with the AC (García-Rosales *et al.*, 2020), behavioural control either by motor commands or volitional vocalization production (Kobler *et al.*, 1987; Eiermann & Esser, 2000; Weineck *et al.*, 2020), and brain-to-brain synchronization during social interactions (Zhang & Yartsev, 2019).

Phase-amplitude coupling in FAF and AC occurs mostly at two conspicuously distinct frequency regimes during spontaneous activity and sound processing (**Figs. 4-6**). We speculate that the different δ/γ_high_ and θ/γ_low_ coupling in frontal and auditory cortices, respectively, indicate that the properties and interactions between the neural substrates at the core of low- and high-frequency oscillations differ across structures. We recently showed that FAF and AC synchronize in low frequencies with and without acoustic stimulation (spanning delta-theta rhythms; (García-Rosales *et al.*, 2020)). In light of the former, one could speculate that PAC further supports the functional relationship between auditory and frontal cortex, bringing together local computations occurring at non-overlapping temporal scales in different gamma sub-bands, according to the synaptic properties of each region. In this case, low-frequency dynamics could provide the temporal basis for fronto-temporal auditory integration. The former would be in accordance with proposed roles of PAC for the facilitation of interareal communication (Colgin *et al.*, 2009; Hyafil *et al.*, 2015b; Helfrich & Knight, 2016). Note, however, that such views remain to be thoroughly addressed in the FAF-AC network. Further research should be aimed at elucidating the synaptic properties of the neuronal networks responsible for delta-theta and gamma oscillations in FAF and AC, and at understanding the function of PAC across regions for higher cognitive demands well beyond passive listening.

### Methodological considerations

The measurements of phase-amplitude coupling from neuronal oscillations can be affected by methodological caveats and several physiological variables (Aru *et al.*, 2015). For example, Tort and colleagues argue that respiratory rhythms, coupled with gamma-band activity, might influence PAC measurements in many cortical areas (Tort *et al.*, 2018). It is nevertheless contended that respiration-related PAC and oscillatory activity are not necessarily artifactual, but that they may reflect cognitive processes and mechanisms for active sensing (Corcoran *et al.*, 2018; Tort *et al.*, 2018). In the particular case of the frontal cortex (where the FAF is located), respiration-related low-frequency oscillations (in the delta-theta bands) modulate a sub-band of gamma with frequencies ranging from 70-120 Hz (Zhong *et al.*, 2017). Interestingly, this falls within the δ/γ_high_ range of this study (note **Figs. 4** and **5**), which makes it possible that the PAC here reported carries signatures of respiration-gamma coupling. As we do not have data from respiratory rhythms recorded simultaneously with the neural activity, the extent of the possible modulation of PAC values by respiration in the FAF cannot be quantified.

We note, however, that if respiration-coupled gamma activity in frontal areas subserve high-order perception (Tort *et al.*, 2010; Corcoran *et al.*, 2018), the PAC associated to respiration may also be important for sensory integration, particularly in the auditory and olfactory modalities. *C. perspicillata* bats rely on multimodal clues for navigation in naturalistic environments, for which olfaction and audition appear crucial (Thies *et al.*, 1998). The interesting possibility of multimodal integration in FAF supported by oscillatory dynamics like PAC, and the extent to which it is modulated by endogenous oscillations that may or may not be related to respiratory rhythms, needs to be thoroughly addressed in future experimental work.

## Materials and Methods

### Animal preparation and surgical procedures

The study was conducted using five awake *Carollia perspicillata* bats (all males). All experimental procedures were in compliance with European regulations on animal experimentation, and approved by the Regierungspräsidium Darmstad (experimental permit #FU-1126). Animals were obtained from a colony at the Goethe-University in Frankfurt. Bats used for experiments were isolated from the main colony.

Surgical and experimental procedures are described in detail in a previous study (García-Rosales *et al.*, 2020), which addressed the functional connectivity between the frontal-auditory field and the auditory cortex this bat species. In brief, for surgery, animals were anesthetized with a mixture of ketamine-xylazine (ketamine: 10 mg*kg^−1^, Ketavet, Pfizer; xylazine: 38 mg*kg^−1^, Rompun, Bayer), and their auditory and frontal cortices exposed by means of a small craniotomy (ca. 1 mm^2^) performed with a scalpel blade. The anatomical location of the two regions of interest was assessed by means of well-described landmarks in both frontal and auditory areas (Esser & Eiermann, 1999; Eiermann & Esser, 2000). After surgery, animals were allowed to recover for at least two days before undergoing experiments. Recordings lasted no more than 4 h per session, and each bat was allowed to recover between sessions for at least a full day. Water was given to the bat at periods of ∼1-1.5 h, and experiments for the day were halted if the animal showed any sign of discomfort.

### Electrophysiological recordings

Electrophysiological data was acquired inside a sound-proof and electrically isolated chamber, where bats were placed on a custom-made holder which was kept at a constant temperature of 30°C with a heating blanket (Harvard, Homeothermic blanket control unit). A speaker (NeoCD 1.0 Ribbon Tweeter; Fountek Electronics, China), used for free-field stimulation, was positioned 12 cm away from the bat’s right ear, contralateral to the hemisphere on which recordings were made. Speaker calibration was done with a ¼-inch microphone (Brüel & Kjær, model 4135, Denmark), connected to a custom-made amplifier.

Recordings were made from the FAF and AC of the left hemisphere. As described in previous studies (Garcia-Rosales *et al.*, 2019; García-Rosales *et al.*, 2020), a NeuroNexus laminar probe (Model A1×16, impedance: 0.5–3 MΩ; 50 μm channel spacing) was carefully inserted perpendicularly into the AC until the uppermost channel was barely visible at the cortical surface. Therefore, the probe’s channels spanned depths from 0-750 μm, covering the extent of an auditory cortical column in *C. perspicillata*’s brain (see (Garcia-Rosales *et al.*, 2019)). Recordings in the FAF were performed, simultaneously to those in the AC, with a single carbon electrode (Carbostar-1, Kation scientific; Impedance at 1 kHz: 0.4–1.2 MΩ), at cortical depths of ∼300-450 μm (313 +- 56 μm; mean +- std).

Both the probe in the AC and the carbon electrode in the FAF were connected each to their own micropreamplifier (MPA 16, Multichannel Systems MCS GmbH, Reutlingen, Germany), which were in turn connected to an integrated amplifier and analog-to-digital converter with 32-channel capacity (Multi Channel Systems MCS GmbH, model ME32 System, Germany). The sampling frequency of the recordings was 20 kHz, and the precision of 16 bits. Data were visualized online and stored in a computer, using the MC_Rack_Software (Multi Channel Systems MCS GmbH, Reutlingen, Germany; version 4.6.2).

### Acoustic stimulation

Acoustic stimulation was controlled with a custom-written Matlab (version 7.9.0.529 (R2009b), MathWorks, Natick, MA) software. Sounds consisted of a natural distress sequence, which is representative of this bat species’ distress repertoire (for recording details and sequence characteristics, see (Hechavarria *et al.*, 2016a; García-Rosales *et al.*, 2018a)), as well as two artificially constructed syllabic trains. One of this trains consisted of a single distress syllable (also representative of *C. perspicillata*’s distress repertoire) repeated isochronously at a rate of 5.28 Hz for a period of 2 s. The second syllabic train consisted of the same syllable repeated with an average rate of 70 Hz, in a Poisson-like manner, with a duration of 4 s. The spectrogram and oscillogram of the sequences, as well as the spectrogram of the syllable used to construct the artificial trains, are depicted in **Fig. 1**.

Auditory stimuli were digital-to-analog converted using a sound card (M2Tech Hi-face DAC, 384 kHz, 32 bit; sampling frequency used: 192 kHz due to technical reasons), amplified (Rotel power amplifier, model RB-1050), and fed to the speaker inside the chamber (description above). Before presentation, sounds were low-pass filtered (80 kHz) and down-sampled to 192 kHz to avoid aliasing artefacts. Stimuli were presented 50 times each, in a pseudorandom order, with an inter-stimulus interval of 1 s. A period of 300 ms and another of 500 ms of silence was padded at the beginning and the end of each sequence, respectively. Before presenting the stimulus battery, 180 s of spontaneous activity were recorded per penetration.

### Separation of local-field potentials

Data analyses were performed offline using custom-written Matlab scripts (version 8.6.0.267246 (R2015b)). The raw signal from each channel (either FAF or the 16 channels in the AC) was band-pass filtered between 0.1 and 300 Hz (4^th^ order Butterworth filter) in order to obtain local-field potentials. For computational reasons, LFPs were down-sampled to 1 kHz and stored in order to be used in subsequent analyses.

### Phase-amplitude coupling

Phase-amplitude coupling was calculated for each stimulus and spontaneous activity per penetration, based on previously published methodology (Kikuchi *et al.*, 2017). For low frequencies, which provided the phase reference, LPFs were filtered (4^th^ order bandpass Butterworth filter) in the following bands: 1-3, 3-5, 5-7, …, and 13-15 Hz, thus having centre frequencies of 2, 4, 6, … 14 Hz, with 2 Hz bandwidth. For higher frequencies, providing the amplitude, LFPs were filtered in bands of 25-35, 30-40, 35-45, …, and 120-130 Hz, therefore having centre frequencies of 30, 35, 40, 45, …, 125 Hz, with 10 Hz bandwidth. After filtering and applying the Hilbert transform on the signals (see below), only the time window between stimulus onset and offset was considered for analysis in a trial.

For analysing sound-related LFP responses, the instantaneous phase [*ϕ(t)*] and amplitude [*A(t)*] of the signal were extracted from low and high frequency filtered LFPs, respectively, by means of a Hilbert transform. To reduce the effect of the stimulus-evoked responses, which could affect PAC values and bias true interactions (Aru *et al.*, 2015), before filtering and determining *ϕ(t)* and *A(t)* per trial, the average of across trials for the current stimulus and penetration (n = 50 trials) was subtracted from the individual response of each trial. The former has the consequence of reducing the effect of time-locked responses in the LFPs for PAC calculations (Kikuchi *et al.*, 2017). Additionally, to minimize the effect of phase non-uniformities (clustering) in the LFPs caused by non-oscillatory periodicities in the field potentials, which could also bias PAC estimates (van Driel *et al.*, 2015), the mean vector of the phase angles was linearly subtracted from the instantaneous phase time series as follows:

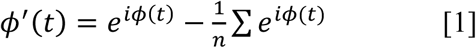

where *ϕ*′(*t*) denotes the corrected phase (i.e. after phase-cluster de-biasing) at time *t*, and *n* represents the number of time points in the series. With *ϕ*′(*t*) and *A(t)*, a composite time series *z*(*t*) = *A*(*t*) * *ϕ*′(*t*) was constructed. From *z(t)* the modulation index (MI) was quantified as:

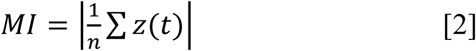

PAC suffers from a number of caveats that depend on the way it is calculated, and on the temporal structure and statistical properties of the LFPs (Aru *et al.*, 2015). As mentioned above, we took measures to minimize possible confounding effects by subtracting the trial-average from each individual trial in response to a stimulus, and by addressing the bias introduced by phase-clustering. In addition to those steps, we calculated a surrogate MI (MI_surr_) by matching the phase series *ϕ*′(*t*) of a given trial *k* with the amplitude *A(t)* of another trial *m* (see **Fig. 3A** and (Aru *et al.*, 2015; Kikuchi *et al.*, 2017)). The trial-shuffling approach further allows to control more stringently for the effect of evoked responses in the LFPs, given that the average evoked-related amplitude and phase responses are unaltered and therefore contribute to the PAC in a similar way as the non-shuffled response. On the other hand, any contribution to the PAC that was trial-by-trial variable would be abolished with the trial-shuffling procedure. Amplitude and phase were paired at random across trials a large number of times (250 permutations), and a modulation index calculated for each iteration. This surrogate method yields a null distribution that accounts for PAC attributable to evoked responses, while allowing to examine trial-specific coupling (Kikuchi *et al.*, 2017).

Modulation indexes obtained with the non-surrogate data (“direct analysis” in **Fig. 3**) were z-normalized to the null distribution obtained by the surrogate approach (zMI). If no effect of PAC exists in the data, zMI values would hover around 0, whereas coupling effects would yield zMIs significantly higher than 0. To assess the former, we used a z-score of 2.5 (i.e. 2.5 standard deviations from the null) as threshold per penetration (see **Supplementary Figure S1**).

The quantification of spontaneous PAC was similar to that of the stimulus-related PAC. LFP chunks (n = 50; same as the number of trials used during stimulation) were selected randomly from the 180 s window of a given penetration, with a length of 1.964 s (the same length as the natural call used for stimulation), and without any overlap. Before chunking, the 180 s window was filtered and Hilbert-transformed for obtaining phase and amplitude series, in order to avoid edge artefacts in the filtered and transformed LFP segments. Because there was no temporal frame of reference, chunk averaging was not performed for subtraction. Chunks were treated as stimulation trials (see above), and PAC analyses together with surrogate calculations were applied in likeness to those performed with stimulus-driven responses.

To test for a population trend of positive PAC, we evaluated, per phase-amplitude frequency pair, whether zMIs for the population were significantly higher than 0 (FDR-corrected, tailed Wilcoxon signed rank test, significance after p_corr_ < 0.05). This is indicated as grey lines in **Figs. 4A** and **5**.

Differences between the PAC in FAF (PAC_FAF_) and AC (PAC_AC_) were calculated by subtracting the PAC maps obtained for the frontal and the auditory cortices, the latter considering 5 representative depths of 50, 300, 450, 600, 750 μm. Significant differences between channels in FAF and AC were determined by means of FDR-corrected Wilcoxon signed rank tests, and significance was considered when p_corr_ < 0.05. Beyond significance testing, we evaluated the effect size of the difference between PAC_FAF_ and PAC_AC_ as follows (Fritz *et al.*, 2012):

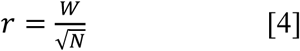

where *r* is the effect size, *W* is the test statistic of the Wilcoxon signed rank test, and N is the sample size (N = 50 penetrations during acoustic processing, and N = 49 during spontaneous activity). Note that the spontaneous activity of one of the penetrations could not be evaluated due to technical reasons. According to (Fritz *et al.*, 2012) values of *r* < 0.3 were considered negligible effects, whereas values of 0.3 ≤ *r* ≤ 0.05 were considered medium, and values of *r* > 0.5 were considered large effects. Only large effects are depicted in the figures as contour lines. Positive large effects (red contour lines) indicate PAC_FAF_ > PAC_AC_, whereas negative large effects (purple contour lines) indicate the opposite.

## Data availability

The data that support the findings of this study are available from the corresponding authors upon reasonable request.

## Author contribution

FGR and JCH designed the study. FGR collected the data, analyzed the data, and wrote the manuscript. FGR, LLJ, EGP, YCC, MK, and JCH discussed the results and reviewed the manuscript.

## Acknowledgements

The German Research Council (DFG) funded this work (Grant No. HE 7478/1-1, to JCH).

