## Supplementary Figure 1 for "Phase-amplitude coupling profiles differ in frontal and auditory cortices"

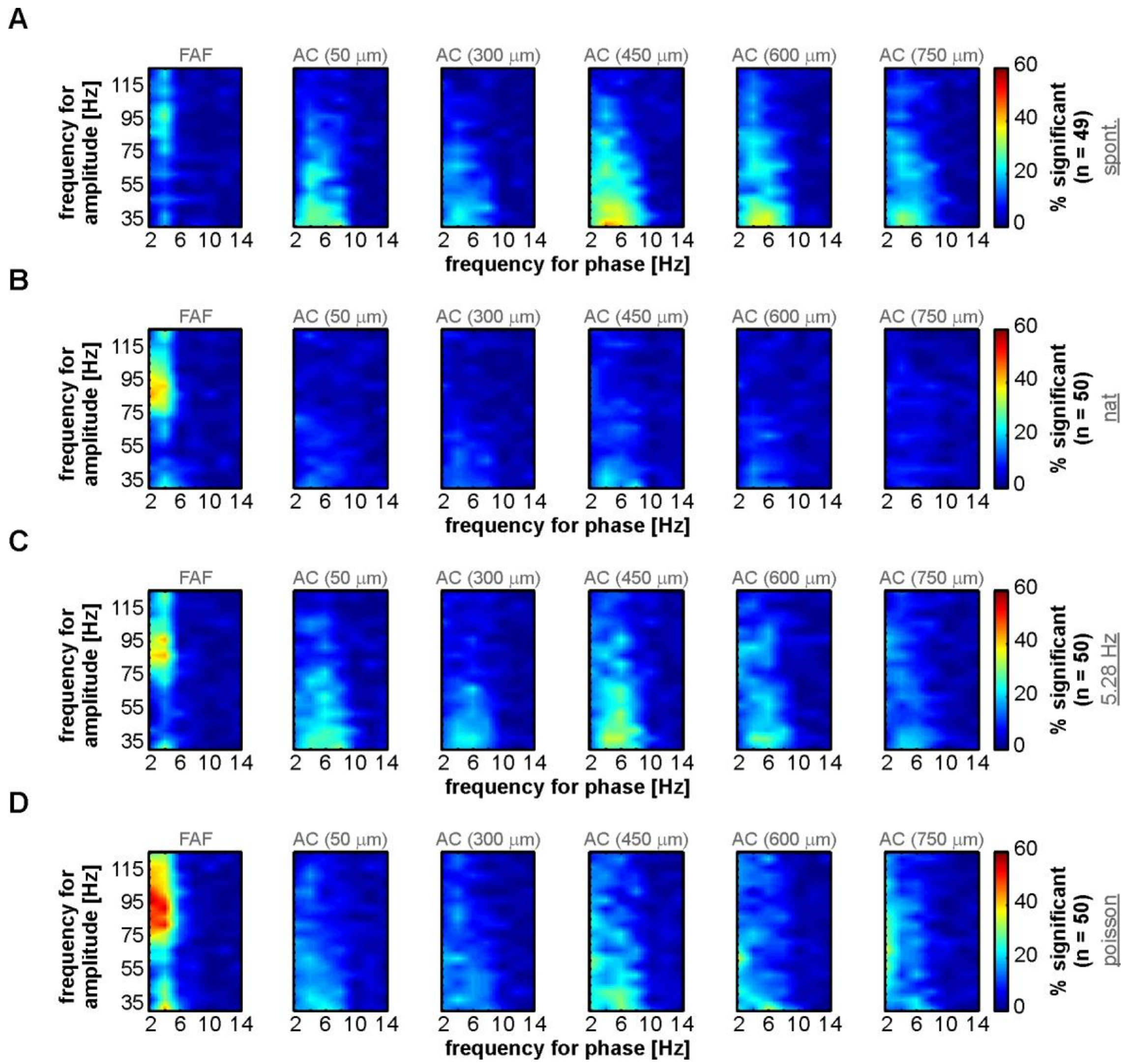


**Supplementary Figure 1. Percentage of significance across penetration in the PAC space.** Each panel shows the percentage of penetrations for which the PAC was significant, at each frequency combination considered (i.e. for phase and amplitude). Panels are organized as follows: **A**, spontaneous activity; **B**, stimulation with the natural call; **C**, stimulation with the 5.28 Hz syllable train; **D**, stimulation with the Poisson syllable train.
